# Transcriptome-based molecular staging of human stem cell-derived retinal organoids uncovers accelerated photoreceptor differentiation by 9-*cis* retinal

**DOI:** 10.1101/733071

**Authors:** Koray D. Kaya, Holly Y. Chen, Matthew J. Brooks, Ryan A. Kelley, Hiroko Shimada, Kunio Nagashima, Natalia de Val, Charles T. Drinnan, Linn Gieser, Kamil Kruczek, Slaven Erceg, Tiansen Li, Dunja Lukovic, Yogita K. Adlakha, Emily Welby, Anand Swaroop

## Abstract

Retinal organoids generated from human pluripotent stem cells exhibit considerable variability in temporal dynamics of differentiation. To assess the maturity of neural retina *in vitro*, we performed transcriptome analyses of developing organoids from human embryonic and induced pluripotent stem cell lines. We show that the developmental variability in organoids was reflected in gene expression profiles and could be evaluated by molecular staging with the human fetal and adult retinal transcriptome data. We also demonstrated that addition of 9-*cis* retinal, instead of widely-used all-*trans* retinoic acid, accelerated rod photoreceptor differentiation in organoid cultures, with higher rhodopsin expression and more mature mitochondrial morphology evident by day 120. Our studies thus provide an objective transcriptome-based modality for determining the differentiation state of retinal organoids, which should facilitate disease modeling and evaluation of therapies *in vitro*.

**Summary Statement:** Three-dimensional organoids derived from human pluripotent stem cells have been extensively applied for investigating organogenesis, modeling diseases and development of therapies. However, substantial variations within organoids pose challenges for comparison among different cultures and studies. We generated transcriptomes of multiple distinct retinal organoids and compared these to human fetal and adult retina gene profiles for molecular staging of differentiation state of the cultures. Our analysis revealed the advantage of using 9-*cis* retinal, instead of the widely-used all-*trans* retinoic acid, in facilitating rod photoreceptor differentiation. Thus, a transcriptome-based comparison can provide an objective method to uncover the maturity of organoid cultures across different lines and in various study platforms.

## INTRODUCTION

Human development requires stringent and coordinated control of gene expression, signaling pathways, and cellular interactions that result in the generation of distinct cell types and tissues with complex morphological and functional phenotypes (1, 2). However, most of our current knowledge of the fundamental molecular events underlying cell-type specification and tissue differentiation has been derived from model organisms. Despite studies using human preimplantation embryos (3, 4) and fetal tissue (5–7), the complexities of human organogenesis are poorly understood (8, 9). Pioneering advances in the generation of human embryonic stem cells (hESCs) (10) and induced pluripotent stem cells (iPSCs) (11), together with the development of three-dimensional (3-D) organoid cultures (12–14), have revolutionized the studies of human development, facilitated individualized disease modeling, and rejuvenated the field of regenerative medicine (15–18).

Retinogenesis begins with specification of the forebrain neuroectoderm, and patterning of the early eye field is governed by finely-tuned regulatory networks of signaling pathways and transcription factors (19). Distinct morphological changes of the rostral neuroectoderm involves lateral expansion of bilateral eye fields to produce the optic vesicles, which invaginate and become the optic cups (20). Landmark studies using fetal and neonatal tissues have provided unique insights, distinct from model organisms, into human retinal development (21–24). By providing appropriate systemic and exogenous cues, human pluripotent stem cells can be directed to self-organize into 3-D optic vesicle or optic cup structures (13, 14). Retinal organoids mimic early eye field development, with VSX2+ (also called CHX10) multipotent retinal progenitor cells differentiating into a polarized and laminated architecture harboring all types of retinal neurons and the Müller glia (25). Rod and cone photoreceptors in organoid culture express opsin and other phototransduction genes (26–28) and develop rudimentary outer segment-like structures at late stages of differentiation (29, 30). However, current methods for characterizing retinal organoids have largely relied upon expression of select cell-type specific markers and histology, which provide limited information about the precise differentiation status. Although live imaging modalities have been employed recently for characterization and developmental staging (31, 32), we still lack molecular insights into retinal organoid differentiation and maturation on a global scale, and how different experimental conditions (e.g., cell lines and/or protocols) could impact organoid cultures.

In this study, we performed transcriptome profiling of developing retinal organoids generated from hESCs and hiPSCs, utilizing modifications of a widely-used protocol (29). Comparative transcriptome analyses with gene profiles of human fetal and adult retina revealed the molecular stages of retinal organoids and demonstrated their differentiation status and cellular composition more accurately. We also identified a specific role of 9-*cis* retinal (9CRAL) in expediting rod photoreceptor differentiation as compared to the currently used all-*trans* retinoic acid (ATRA). Thus, our studies establish a transcriptome-based molecular staging system using human fetal and adult data, enabling direct comparison of organoids under different experimental conditions for disease modeling and evaluation of therapies.

## MATERIALS AND METHODS

### Maintenance and differentiation of human pluripotent stem cells

CRX-GFP H9 is a subclone of H9 human embryonic stem cell (ESC) line, carrying a green fluorescent protein (GFP) gene under the control of the cone-rod homeobox (*CRX*) promoter as previously reported (26). Human induced pluripotent stem cell (iPSC) line PEN8E and NEI377 were reprogrammed from skin biopsies using integration-free Sendai virus carrying the four Yamanaka factors, as described (33), and their genome integrity and pluripotency have been evaluated (Shimada et al., 2017). ESP1 and ESP2 lines were reprogrammed by Oct4, Klf4, Sox2, c-Myc, Lin-28 mRNAs (34) and Sendai virus (Ctrl1 FiPS4F1, Spanish National Stem Cell Bank), respectively. H9, PEN8E and NEI377 were maintained in Essential 8 medium (E8; ThermoFisher Scientific) under hypoxia (5% O_2_), and ESP1 and ESP2 lines in mTeSR1^®^ (Stem Cell Technologies) under normoxia (20% O_2_). All lines were sustained on BD Matrigel™ human embryonic stem cell-qualified Basement Membrane Matrix (Corning)-coated plates. PSCs were passaged every 3-4 days at 70-80% confluency using the EDTA-based protocol (35).

To start differentiation, all lines were detached and dissociated into small clumps with the EDTA dissociation protocol (35) and cultured in polyHEMA-coated or Ultra Low Attachment Culture Dishes (Corning) in E8 with 10 μM Y-27632 (Tocris) to form embryoid bodies (EBs). Neural-induction medium (NIM) consisting of DMEM/F-12 (1:1), 1% N2 supplement (ThermoFisher Scientific), 1x MEM non-essential amino acids (NEAA), and 2 μg/ml heparin (Millipore Sigma) was added at differentiation day (D)1 and D2 to reach a final ratio of 3:1 and 1:1 NIM:E8, respectively. Starting at D3, EBs were cultured in 100% NIM for an additional 4 days. EBs were collected at D7 and plated onto BD Matrigel™ Growth Factor Reduced (GFR) Basement Membrane Matrix (Corning)-coated plates. On D16, medium was changed to photoreceptor induction medium (PIM) consisting of DMEM/F-12 (3:1) with 2% B27 without Vitamin A, 1% antibiotic-antimycotic solution (ThermoFisher Scientific), and 1X NEAA. Medium was changed daily.

Upon appearance of optic vesicles (OVs; typically, D21-28) with neuroepithelium morphology, the regions were excised with tungsten needles under magnification and transferred to suspension culture in polyHEMA-coated or Ultra Low Attachment 10-cm^2^ Culture Dishes (Corning) in PIM. After D42, PIM was supplemented with 10% fetal bovine serum (FBS; ThermoFisher Scientific), 100 mM Taurine (Millipore Sigma), 2 mM GlutaMAX (ThermoFisher Scientific). Starting at D63, PIM was supplemented with 1 mM retinoid (Millipore Sigma) three times a week and switched to 0.5 mM at D92. 1% N2 supplement was included starting at D92. Medium for H9 and PEN8E-derived organoids was supplemented with 20 ng/ml insulin-like growth factor 1 (IGF1; ThermoFisher Scientific) and 55 nM beta-mercaptoethanol (2-ME; ThermoFisher Scientific) from dissection till the end of differentiation. 9-*cis* retinal (9CRAL; Millipore Sigma) was added at the described concentrations and corresponding time period during media change. Organoids for dataset PEN8E_2 were cultured with 10 ng/ml IGF1 from D42 to D200 and all-*trans* retinoic acid (ATRA; Millipore Sigma) was supplemented to PIM at 1 μM 5 times per week and at 0.5 μM twice a week for D63-92 and D92-180, respectively. ESP-derived organoid cultures were maintained in the same manner as PEN8E_2 organoids, except no IGF1 was used. For all experiments directly comparing 9CRAL and ATRA, media changes were performed in the dark under dim red light to reduce isomerization of the retinoids. Retinal organoids were collected in the dark and processed in the light as described for all other experiments.

#### Immunofluorescence

hPSC-derived retinal organoids were collected throughout the differentiation process, fixed, and immunostained as previously described (36). Briefly, the organoids were fixed in 2% or 4% paraformaldehyde in 1X PBS for 1 hour at room temperature. Fixation was followed by cryoprotection in a sucrose gradient, 10-30%. Retinal tissues were then embedded in Shandon M1 embedding matrix and frozen on dry ice. 10μm sections were obtained using a Leica cryostat at −14°C. Sections were placed on super frost plus slides and stored at −20°C until use. Specific antibodies and concentrations are summarized in Table S5.

#### Immunoblots

The protocol for immunoblot analysis is as previously described (37). Individual organoids were lysed in 50 μL of 1% triton X-100 (Millipore Sigma) in 1X phosphate-buffered saline (PBS; ThermoFisher Scientific) supplemented with 1X protease inhibitors (Roche). Gentle trituration was then used to dissociate the organoid until a homogenous solution was obtained. Samples were then incubated on a nutator at 4°C for 1 hour and centrifuged at 4°C at 4000xg for 5 minutes. Protein concentration was measured via Pierce BCA protein assay (ThermoFisher). Approximately 15 μg supernatant protein was diluted 4:1 in reducing 4X Laemmeli buffer for 1 hour at room temperature and separated at 100V for 1 hour on 10% SDS-PAGE gels, which were then transferred to PVDF membranes at 100V for 1 hour. After blocking in 5% milk in 1X TBST (ChemCruz with 1mM EDTA, Santa Cruz Biotech) for 1 hour at room temperature, blots were incubated in rhodopsin antibody cocktail (1:1000 1D4, 3A6, and 4D2, generous gift from Dr. Robert Molday) overnight in 1% milk in 1X TBST. The following morning membranes were washed in 1X TBST shaking at room temperature four times for 10 minutes each. Membranes were then incubated in appropriate secondary species IgG conjugated to horseradish peroxidase (1:10000) in 1X TBST shaking at room temperature for 1.5 hours. Membranes were then washed again four times in 1X TBST for 10 minutes each. Prior to imaging, the membranes were exposed to supersignal^®^ west pico ECL solution (ThermoFisher) for 5 minutes and chemiluminescence was captured using a Bio-Rad ChemiDoc™ touch (BioRad). Membranes were next dried overnight at room temperature to remove antibody binding to protein. Dry membranes were reactivated in 100% methanol and incubated with gamma-tubulin (1:1000, ab11317, Abcam) overnight as described above. Images were analyzed in Image Lab (BioRad) and exported to Adobe photoshop for figure generation.

#### Transmission electron microscopy

Retinal organoids were processed for EM analysis as previously described (33). Briefly, the organoids were initially fixed in 4% formaldehyde and 2% glutaraldehyde in 0.1M cacodylate buffer (pH 7.4) (Tousimis) for 2 hours, washed in cacodylate buffer 3 times, then fixed in osmium tetroxide (1% v/v in 0.1M cacodylate buffer) (Electron Microscope Science) for 1 hour in a room temperature. The organoids were washed in the same buffer 3 times, followed by acetate buffer (0.1M pH 4.2), and *en-bloc* staining in uranyl acetate (0.5% w/v) (Electron Microscope Science) in acetate buffer for 1 hour. The samples were dehydrated in ethanol solution (35%, 50%, 75%, 95% and 100%), followed by propylene oxide. Subsequently, the samples were infiltrated in a mixture of propylene oxide and epoxy resin (1:1) overnight, embedded in a pure epoxy resin in a flat mold, and cured in 55°C oven for 48 hours. Thin-sections (70 to 80nm) were made with an ultramicrotome (UC 7) and diamond knife (Diatome), attached on 200-mesh copper grid, and counter stained in aqueous solution of uranyl acetate (0.5% w/v) followed by lead citrate solutions. The thin sections were stabilized by carbon evaporation under a vacuum evaporator prior to the EM examination. The digital images were taken in the electron microscope (H7650) equipped digital camera (AMT).

#### RNA-seq Analysis

High quality total RNA (100 ng, RIN > 7) from at least two independent replicates for hPSCs at various differentiation time points were subjected to mRNA directional library construction as described previously (26, 38). Paired-end sequencing was performed to a length of 125 bases using HiSeq2500 (Illumina, San Diego, CA). Genome reference sequence GRCm38.p7 and Ensembl v82 annotation was used for alignment and quantitation. Quality control, sequence alignment, transcript and gene-level quantification of primary RNA-seq data were accomplished using an established bioinformatics pipeline (39). Gene expression clustering was performed on gene-wise Z-scores using Affinity Propagation (AP) with apcluster v1.4.7 package (40, 41) in R statistical environment. GO enrichment analysis was performed using clusterProfiler v3.6.0 (42). Dynamic time warping (DTW) analysis was performed using the dtw v1.20-1 package in R. CPM (log_2_) values from the retina-centric gene set (Table S3) were used to generate the Local Cost Matrix. Co-inertia analysis was performed using MADE4 v1.52.0 package in R. Transcriptome of adult human retina samples was obtained from NEI Commons (Brooks, MJ and Swaroop, A, https://neicommons.nei.nih.gov).

#### Alignment and transcript quantitation

Data generated for the five different cell lines transcriptome comparison and 9CRAL/ATRA analysis (used in Figure 4) were analyzed separately. For each dataset, genes were kept for further analysis only if there were ≥5 count per millions (CPM) in all replicates of at least one group of the datasets. The data were subjected to TMM normalization by edgeR v3.20.9 (43, 44), and PCA and Pearson correlation were performed with normalized CPM (log_2_) values. Differential expression analysis was performed using limma v3.34.9 (45), and genes having ≥ 2-fold change at least between 2 time points in either transcriptome of comparison and a false discovery rate (FDR) ≤ 0.01 were considered to be significantly differentially expressed.

#### Cluster analysis

Gene expression clustering was performed on gene-wise Z-scores using Affinity Propagation (AP) (apcluster v1.4.7 package) (40, 41) with k-means option where k=12 by choosing corSimMat function as similarity measure. Determination of the cluster number k was accomplished by empirical observation of the heatmap produced from the PCA projections of genes-wise z-score.

#### Gene ontology analysis

GO enrichment analysis was performed using clusterProfiler v3.6.0 (42). To reduce redundancy associated with GO analysis, we performed Wang semantic similarity comparisons of the enriched GO terms using GOSemSim v2.4.1 (46) and semantic similarities clustered with AP. The term with the highest level in each cluster is taken as the enriched GO term. Heatmaps of GO term gene lists were generated using the log2 CPM values. The GO enrichment ratio is calculated with reference to the number of pathway genes found overlapping with the analysis.

#### Time-course stage comparison of retinal organoids with human fetal retina

Open-Ended Dynamic Time Warping (OE-DTW) analysis was used to compare the maturation state of the different retina organoid time-course stage expression data to human fetal retina (GSE104827), human adult retina, as previously described (47). Retina organoid time-course stages were defined as: Stage 1: 25-36 days, Stage 2: 50-75 days, Stage 3: 80-90 days, Stage 4: 105-125 days, Stage 5: 145-172 days, and Stage 6: 186-205. Mean gene expression values at each stage were used for OE-DTW analysis. The gene set used for OE-DTW analysis consisted of 118 well-known retina-centric transcription factors and cell-type markers (see Figure S4B, Table S3).

### RESULTS

#### Comparable morphology between hESC and hiPSC-derived retinal organoids

Two human pluripotent stem cell lines, H9 (hESC) and PEN8E (hiPSC), were differentiated into retinal organoids using a widely-used robust protocol (29), with minor modifications based on our mouse organoid differentiation method that included 9CRAL (36). Both lines could generate phase-bright neural retina with cone and rod photoreceptors of comparable morphology (Figure 1A-C). S opsin (OPN1SW) was polarized to the apical side of organoids (outer surface of organoid, exposed to media) as early as Differentiation Day (D) 90, with further increase in expression from D120 to D200. L/M cone photoreceptors (labeled with L/M opsin, OPN1L/MW) were barely evident at D120 but increased significantly at D200 (Figure 1C, left). Rhodopsin (RHO) immunostaining was observed at the apical side as early as D120 in both H9- and PEN8E-derived organoids and elongated as the photoreceptors matured (Figure 1C, right). Retinal ganglion cells (RGCs) were evident at early stages but could not be maintained through the end of differentiation, while all other neural retina cell types showed similar morphology and development in both H9- and PEN8E-organoids (Figure S1).

**Figure 1:**
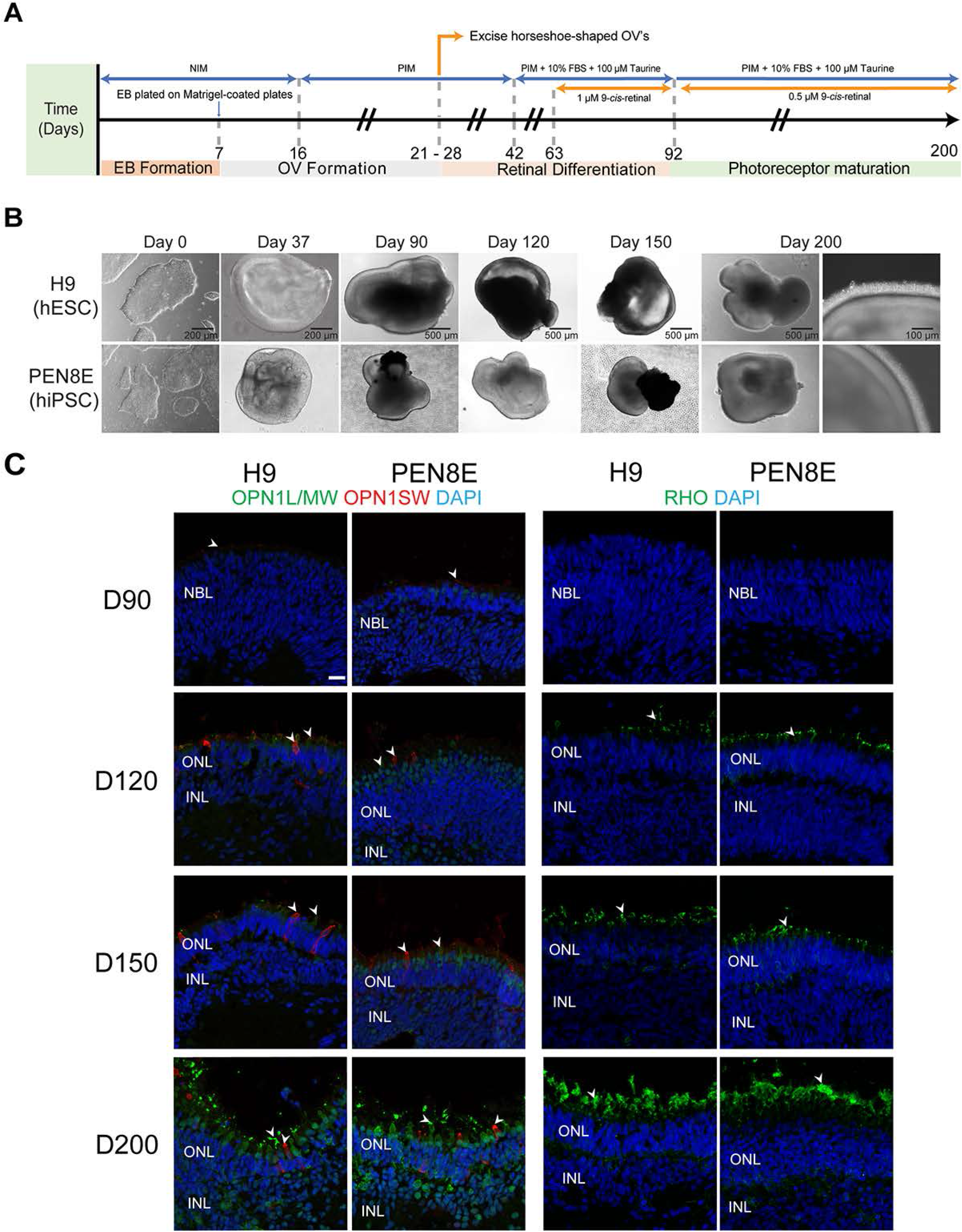
**(A)** Differentiation protocol used in this study, modified from (29, 33). Numbers under the arrow indicate the differentiation day. NIM: neural induction medium; PIM: photoreceptor induction medium; EB: embryoid bodies; OV: optic vesicles; FBS: fetal bovine serum. **(B)** Representative brightfield images of human embryonic stem cells (hESC; H9) and induced pluripotent stem cells (hiPSC; PEN8E), and of differentiating organoids (from Day 37 to Day 200). **(C)** Immunohistochemistry analysis of H9 and PEN8E-derived retinal organoids using marker antibodies for cones (OPN1L/MW, OPN1SW) and rods (RHO). Nuclei were stained with 4’, 6-diamidino-2-phenylindole (DAPI, blue). Arrowheads indicate relevant staining of each marker. Scale bar: 20 μm. D: differentiation day; NBL: neuroblastic layer; ONL: outer nuclear layer; INL, inner nuclear layer; OPL: outer plexiform layer; IPL: inner plexiform layer.

#### Differentially-expressed (DE) gene clusters during organoid development

We then performed RNA-seq analysis of developing retinal organoids to decipher major differentiation stages *in vitro*. H9 and PEN8E-derived retinal organoids showed a substantial overlap of a total of 3,851 differentially expressed (DE) genes that exhibited significant changes in expression during differentiation (Figure 2A, Table S1,2). The 1041 DE genes unique to H9 and the 1749 genes unique to PEN8E showed the same general trend in both datasets (Figure S2A). We noted that many of these genes were significantly differentially expressed in both datasets under a less stringent cutoff of 1.5-fold and 5% FDR (data not shown). Hierarchical clustering analysis of the common DE genes between two databases yielded 8 clusters (C) with either monotonically increasing or decreasing expression (Figure 2B, Table S2B). Gene Ontology (GO) Biological Process enrichment analysis uncovered key biological pathways and genes that could be associated with specific stages of retinal development and identified significantly enriched terms for all clusters except C1 and C7 (Figure 2C). C2-C4 genes displayed a gradual decrease throughout differentiation and were associated with negative regulation of neuronal differentiation, mitotic cell division, and neurodevelopmental processes and signaling (Figure 2D, Figure S3). C4 also included axon guidance genes, which likely reflected the loss of ganglion cells in the organoid cultures. In contrast, C5, C6, and C8 genes exhibited progressively increasing expression during organoid development and included genes associated with neuronal/retinal differentiation and functions including visual perception, synaptogenesis, and phototransduction.

**Figure 2:**
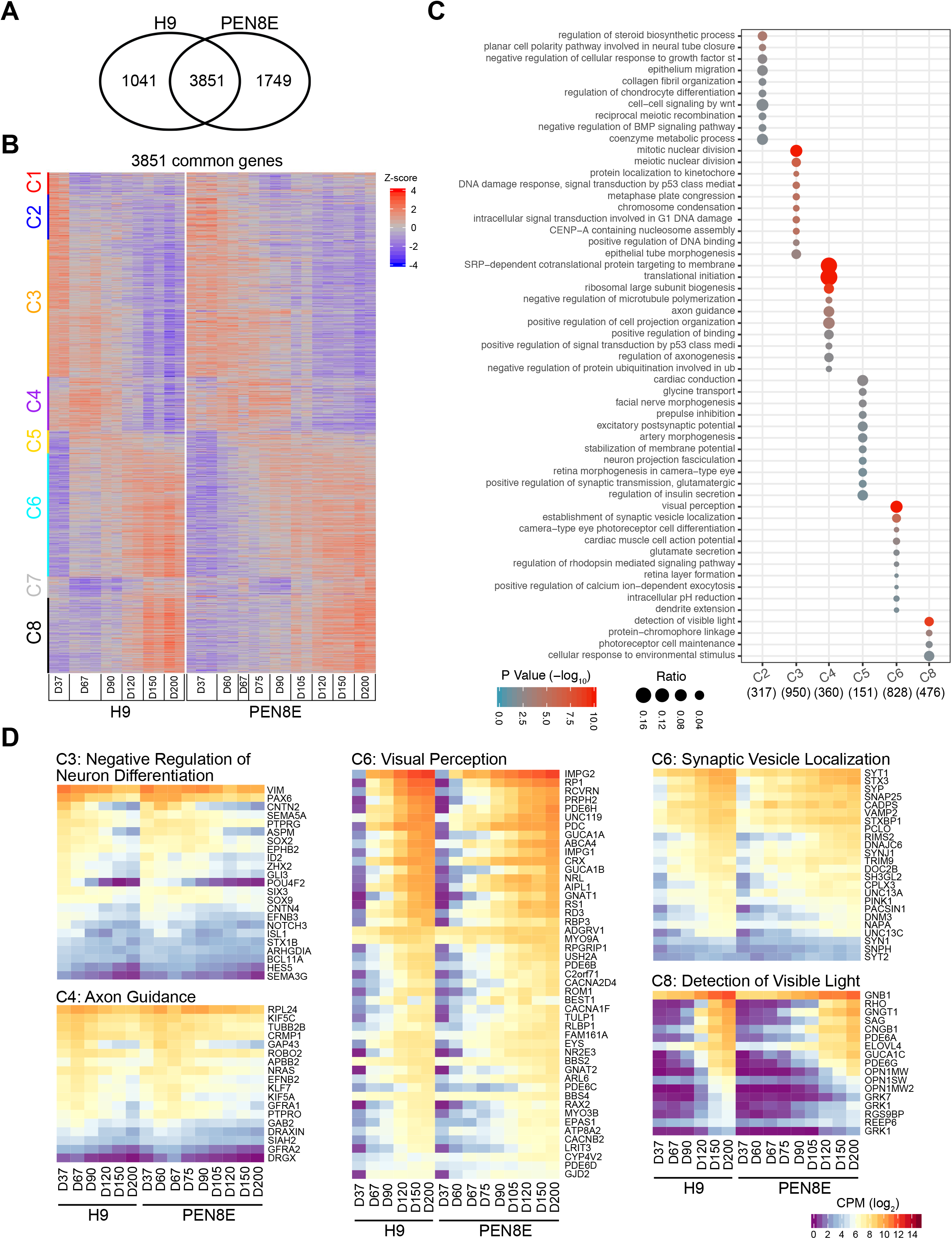
Comparative transcriptome analysis of H9 (hESC) and PEN8E (hiPSC)-derived retinal organoids. **(A)** Venn diagram showing differentially expressed (DE) genes during organoid development from H9 and PEN8E lines. A vast majority of DE genes are common (3851), with 1041 and 1749 unique to H9 and PEN8E, respectively. **(B)** Heatmap of 3851 common DE genes, sorted and clustered based on row-wise z-score expression values in H9. Eight clusters are evident. **(C)** Reduced-redundancy GO Biological Process enrichment analysis of genes in each cluster. Cluster 1 and 7 did not detect any significantly enriched biological processes. **(D)** Heatmaps of GO cluster related genes and their expression patterns in H9 and PEN8E retinal organoids. Expression CPM (log_2_) values were used for analysis; color scale shown at bottom right.

#### Transcriptomes of organoids from different iPSCs and protocols

To evaluate the impact of iPSC lines and/or modifications in the differentiation protocol, we compared the expression of 3,851 DE genes in RNA-seq data of organoids produced from different hiPSC lines (ESP1, ESP2, and NEI377) and/or protocols (PEN8E_2) (see Supplementary Experimental Procedures for details). Principle Component Analysis (PCA) revealed a similar temporal progression in all organoid transcriptomes with increment in differentiation days, as evident from PC1 (Figure S4A). As the harvesting time points varied among individuals and laboratories, we organized the organoid samples into 6 groups, for ease of comparison, based on the collection day (chronological age) and state of differentiation (Figure 3A) and performed PCA (Figure 3B) using a selection of established, retina-centric genes (Table S3). While all organoids clustered together at an early stage of differentiation (group 1), H9- and PEN8E-derived organoids (that included 9CRAL supplementation) showed accelerated differentiation from group 3 and onward, compared to ESP, NEI377, and PEN8E_2 organoids (that used ATRA as described by (29)). In concordance, ESP, NEI377, and PEN8E_2 organoids displayed lower expression of genes associated with phototransduction and outer segments compared to H9- and PEN8E-derived organoids (Figure S4B). NEI377 and PEN8E_2 organoids were generated by an identical protocol and they displayed a similar developmental pattern, except for group 6 of PEN8E_2, which is likely due to iPSC line differences.

**Figure 3:**
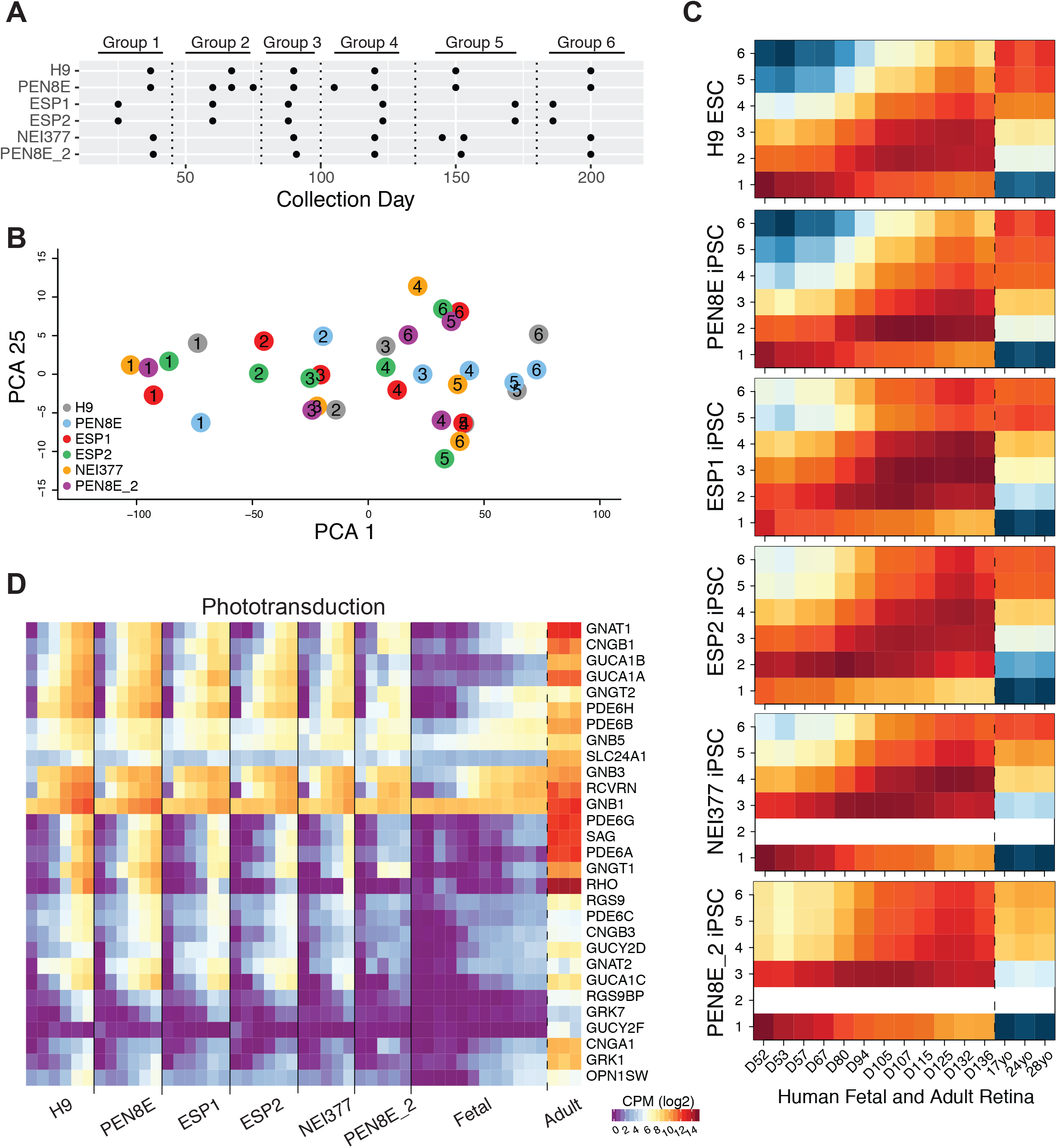
Comparative transcriptome analysis of hPSC-derived organoids and human retinal samples. **(A)** Table showing organoid collection groups (Group 1: D25-37, Group 2: D50-75, Group 3: D80-90, Group 4: D105-125, Group 5: D145-172, Group 6: D185-200). **(B)** PCA plot of grouped hPSC-derived organoid samples based on retinal gene expression. PC1 shows the highest variation percentage and relates to time progression, while PC25 shows an insignificant principle component used for ease of visualization. **(C)** Dynamic Time Warping analysis showing the LCM for all hPSC-derived organoid groups and human fetal (D52-136) and adult (17, 24, 28 years) samples. Warmer colors indicate lower levels of global dissimilarity (lowest: red, medium: orange), whereas cold colors (yellow and blue) represent higher levels of dissimilarity between samples. **(D)** Heatmap demonstrating Mean expression CPM (log_2_) value expression profiles of phototransduction across all hPSC-derived organoids and human retinal samples.

#### Molecular staging of retinal organoids based on developing human retina *in vivo*

In order to determine the maturity of organoid cultures using objective parameters, we performed open-ended dynamic time warping (DTW) analysis to align time series expression data between the organoids and human fetal and adult retina gene profiles (Figure 3C) using a set of retina-centric genes (Table S3). Across all stem cell lines, group 1 organoids (Day 25-37) matched with the earliest human fetal time points (Day 52-67). Subsequently, distinct PSC organoids showed differences in their highest correlation to corresponding human retinal stages. While organoids derived from ESP1, ESP2, NEI377, and PEN8E_2 lines revealed higher correlation with fetal retina samples; groups 4-6 H9 and PEN8E organoids were more concordant with late fetal and adult retina (Figure 3C). For example, group 4 H9 and PEN8E-derived organoids (Day 145-172) were highly correlated with adult samples (17, 24, 28 years of age), whereas ESP1, ESP2, NEI377, and PEN8E_2 organoids corresponded with late fetal samples (Day 105-136) at a similar stage (Figure 3C).

To confirm the result of DTW analysis, we compared the expression of mature retinal genes associated with phototransduction and synaptic function between organoids and *in vivo* retina. Late stage (group 5 and 6) H9 and PEN8E organoids showed similar expression of phototransduction genes as that of the adult retina (Figure 3D). However, most genes from “Transmission across chemical synapses” pathways were down-regulated in organoid cultures, suggesting the retinal organoids were not yet fully mature (Figure S4C).

#### Accelerated rod photoreceptor differentiation by 9CRAL

Much like its physiologically relevant isomer 11-*cis* retinal, 9CRAL can also bind opsin to form functioning rhodopsin, and RA can be generated as an oxidation product. We therefore hypothesized that replacing RA with 9CRAL in our modified protocol would result in improved rod differentiation. To evaluate this hypothesis, we performed a direct comparison of D90 and D120 H9 organoids cultured with ATRA or 9CRAL from D64 onwards. We then applied co-inertia analysis to the transcriptomes of ATRA and 9CRAL-supplemented organoids and developing human fetal retina (D53 to D136). The D90 and D120 of 9CRAL organoid transcriptomes corresponded with the D80-D94 and D125-D136 human fetal retina gene profiles, respectively. However, D90 and D120 ATRA transcriptomes matched with earlier fetal retina time points (D57-67 and D105-115, respectively) (Fig. 4A). Thus, treatment with 9CRAL expedited retinal development in organoid cultures and showed closer temporal transcriptome dynamics to human fetal retina. DE genes between D90 and D120 in ATRA (1284 genes) and 9CRAL (852 genes) data presented an overlap of 594 genes (Figure 4B, Table S4), which could in turn be grouped into 5 clusters by hierarchical clustering analysis (Figure 4C). Redundancy-reduced Gene Ontology (GO) analysis of common DE genes revealed monotonic decrease of C1 and C2 genes involved in cell cycle, suggesting neural progenitor cells exit cell cycle and become mature in retinal organoids (Figure 4D). C3-C5 genes were involved in energy metabolism and visual perception, which further characterized their progressive increase from D90 to D120. Although ATRA and 9CRAL organoids showed comparable developmental patterns, changes of DE genes during development varied between the two groups (Figure 4C). Day-matched comparison of DE genes revealed lower expression of genes involved in regulation of retinoic acid (RA) signaling pathway, RA metabolic processing and energy metabolism, whereas higher expression of visual perception and function genes was evident in both D90 and D120 9CRAL-treated organoids (Figure 4E). Heatmap of early eye field transcription factors and photoreceptor genes consistently showed an expedited photoreceptor differentiation in 9CRAL organoids, as revealed by higher expression of these genes, compared to ATRA organoids (Figure 4F).

**Figure 4:**
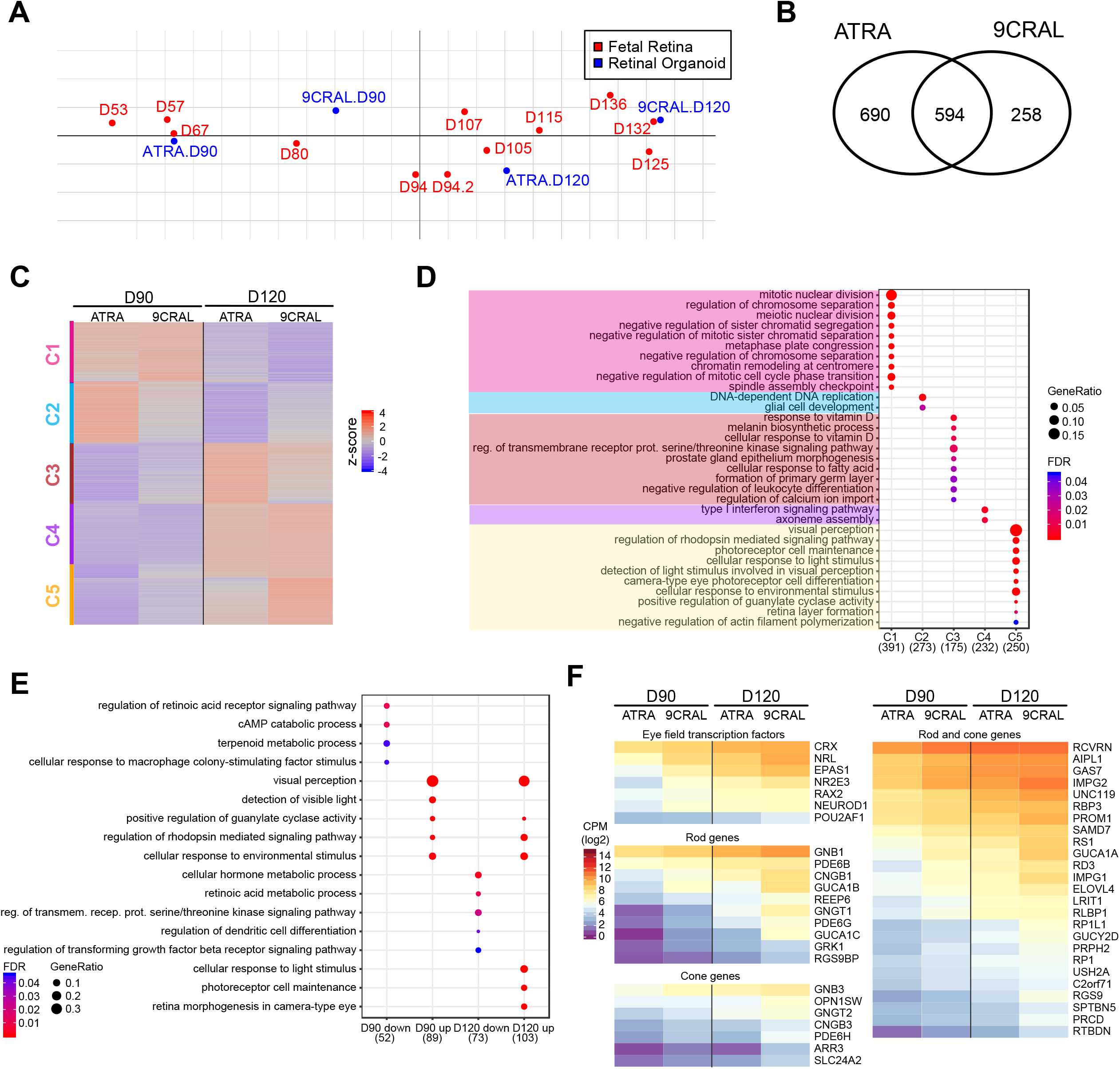
Transcriptome analysis of D90 and D120 organoids treated with all-*trans* retinoic acid (ATRA) or 9-*cis* retinal (9CRAL). **(A)** Co-inertia analysis projecting ordinations of maximum covariation of D90 and D120 ATRA and 9CRAL organoids with human fetal transcriptome data. **(B)** Venn diagram revealing differentially expressed (DE) genes during differentiation between ATRA and 9CRAL organoids. **(C)** Heatmap of 594 common DE genes, sorted and clustered based on row-wise z-score expression values in H9. Five clusters are evident. **(D)** Reduced-redundant GO Biological Process enrichment analysis of genes in each cluster. **(E)** GO analysis of DE genes comparing 9CRAL to ATRA at D90 and D120 (daywise comparison). **(F)** Heatmaps showing expression profiles of genes encoding eye field transcription factors and photoreceptor genes.

To validate our findings that 9CRAL accelerated photoreceptor differentiation, we performed immunohistochemistry on D90 and D120 organoids. D90 9CRAL organoids showed lower expression of retinal progenitor cell marker VSX2 (also called CHX10) and higher expression of a pan-photoreceptor marker (recoverin, RCVRN) compared to ATRA organoids (Figure 5A, upper panel). At D120, 9CRAL-treated organoids showed distinctively more RHO+ cells, which were barely detected in ATRA organoids (Figure 5A, bottom panel), consistent with the higher rhodopsin expression in 9CRAL (14.0±9.0 CPM) compared to ATRA (3.8±1.7 CPM) RNA-seq data. No marked difference was observed in the expression or morphology of L/M-opsin and S-opsin+ cells (see Figure 5A, bottom panel), as well as other neural retina cell types (Figure S5). In concordance, immunoblot analysis demonstrated increased rhodopsin expression in 9CRAL organoids compared to ATRA organoids at D120 (Figure 5B). Ultrastructural analysis of D130 retinal organoids using transmission electron microscopy (TEM) showed an advanced stage of maturation of 9CRAL photoreceptors compared to ATRA (Figure 5C). Ellipsoid (apical) side of photoreceptor inner segments in 9CRAL organoids contained a higher number of mitochondria with typical morphology, which could explain the divergent expression of genes involved in energy metabolism between 9CRAL and ATRA organoids. No significant differences were evident in the morphology of photoreceptor connecting cilia between the two groups. Taken together, our data demonstrates that 9CRAL supplementation expedited differentiation and maturation of rod photoreceptors in retinal organoids.

**Figure 5:**
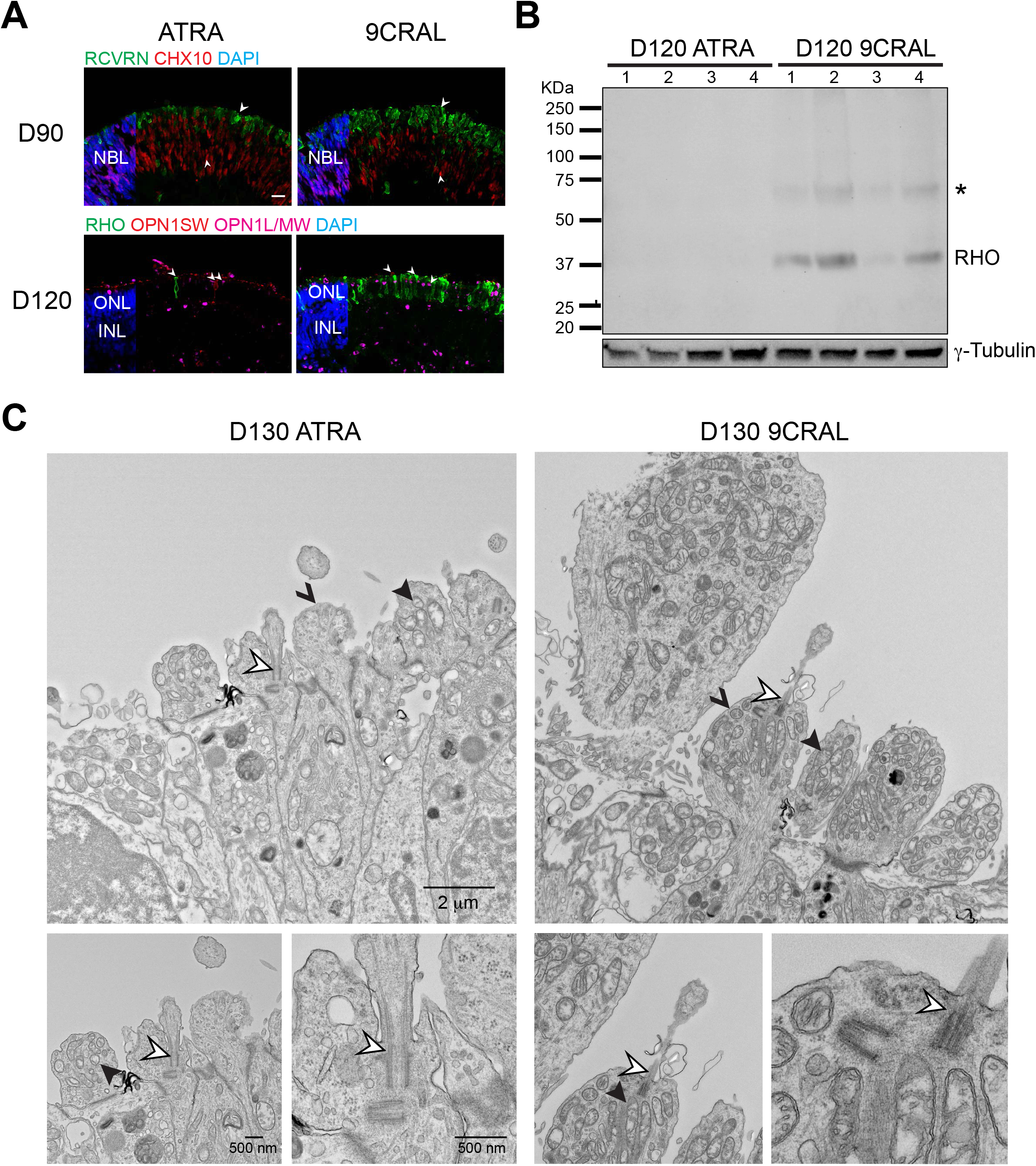
9-*cis* retinal (9CRAL) expedited photoreceptor development. **(A)** Representative images of immunostained sections of H9 retinal organoids supplemented with either ATRA (left) or 9CRAL (right). D90 (top) 10μm sections were immuno-labeled for pan-photoreceptor marker recoverin (green) and retinal progenitor cell marker CHX10 (red). D120 (bottom) were immuno-labeled for rod photoreceptor marker rhodopsin (green), cone photoreceptor marker OPN1SW (red), and L/M cone photoreceptor marker OPN1L/MW (magenta). Nuclei were stained with 4’, 6-diamidino-2-phenylindole (DAPI, blue). Arrowheads indicate relevant staining of a specific marker. Scale bar, 10μm. **(B)** Immunoblot showing rhodopsin expression in H9 ATRA and 9CRAL organoids at D120 (individual replicates are shown). The asterisk denotes a second band which is likely to be dimerization of rhodopsin. γ-tubulin is included as the protein loading control for protein amounts. **(C)** Transmission electron microscopy of longitudinal sections at D130, showing inner segments and cilia. Hollow, solid, and v-shaped arrowheads indicate relevant structure of photoreceptor cilium, mitochondria and inner segments, respectively.

### DISCUSSION

The generation of *in vitro* 3-D organoids from pluripotent stem cells has permitted rapid advances in our understanding of human organogenesis, disease mechanisms, and therapeutic interventions (16, 18). Relatively easy access and promising applications of cell replacement therapy to alleviate blinding diseases have prompted an explosion of studies on human retinal organoids, which exhibit appropriate stratified architecture and differentiation of all relevant cell types (26, 29–31, 48–51). However, lack of appropriate photoreceptor outer segments and synaptic connectivity, and loss of RGCs after prolonged cultures have hampered the progress. The variability in temporal differentiation, depending on protocols and iPSC lines, also demands objective criteria for staging of human retinal organoids. Live imaging and reporter quantification assays have been used to characterize organoid development and staging (31, 52). More recently, retinal organoids from 16 hPSC lines were examined using multiple structural criteria that were then employed for developing a staging system (32). In comparison, our molecular staging provides a more objective measure of organoid maturity based on molecular staging with human retinal development. Our molecular staging profiles are concordant to the recent report describing similar temporal gene expression between *in vivo* human retina and *in vitro* cone-rich retinal organoid transcriptome data (53).

Human iPSCs harbor epigenetic memory of their somatic tissue of origin, which appears to favor their subsequent differentiation towards the lineage related to the donor cells and restrict other cell fates (54). The presence of epigenetic signatures of donor tissues depends on reprogramming methods and passages of iPSCs (55); therefore, differentiation capacity and therapeutic potential of hESCs and hiPSCs are not easily comparable. Interneurons in mouse retinal organoids differentiated from iPSCs cannot be well differentiated due to epigenetic markers of fibroblasts (5) but this phenotype could be alleviated in cultures with additional nutrients (36, 39), suggesting that culture conditions as well as cell lines impact the epigenome in iPSCs. In this report, we differentiated genetically unmatched hESC and hiPSC line reprogrammed from fibroblasts into retinal organoids, and our data showed a similar differentiation capacity of hESC and hiPSC in retinal lineages. Our results are in agreement with a recent study showing equivalent gene expression and neuronal differentiation potential between hESCs and iPSCs (56).

Human organoid cultures manifest substantial variability that may result from multiple factors including intrinsic genetic variations and epigenome state of the iPSC lines, reprogramming method and differentiation protocol. A recent report used a TaqMan array-based analysis of key marker genes to demonstrate differences in retinal organoids (57). Our comprehensive transcriptome-based molecular staging method utilizes global gene expression profiles and can robustly capture organoid variability and developmental status by comparing these to *in vivo* retinal transcriptome data. The gene profiles of ESP1, ESP2, NEI377, and PEN8E_2 showed broadly similar gene profiles and developmental trajectories, suggesting a minimal impact of cell-line intrinsic and protocol variations. On the other hand, H9 and PEN8E clearly exhibited a more mature developmental trajectory and transcriptomes, resulting from the use of 9CRAL in the organoid cultures.

The studies reported here demonstrate accelerated rod photoreceptor differentiation and detection of rhodopsin protein by immunoblot analysis in 9CRAL-supplemented organoids as early as D120. We suggest that 9CRAL is driving more cells towards a photoreceptor cell fate. In our experimental protocol, all media changes were performed in the dark under dim red light to avoid isomerization of 9CRAL to all-*trans* retinal; therefore, at least some of the 9CRAL is likely oxidized into 9-*cis* retinoic acid (9CRA) inside the cells. 9CRA is a potent agonist for both retinoid X receptors (RXRs) and retinoic acid receptors (RARs), while ATRA binds only to RARs (58–60). Retinoic acid is shown to promote photoreceptor development (61–63) and induce the expression of rod differentiation factor NRL (64). We should note that retinoid-related orphan receptor beta (RORβ) regulates rod development by activating NRL (65). In general, the interplay of retinoic acid receptors with other nuclear receptors has substantial impact on transcriptional regulation of genes involved in photoreceptor development (66). Thus, expedited rod differentiation by 9CRAL could result from more potent activation of retinoic acid receptors that may induce rod genes through NRL-regulated gene network (67). Nevertheless, further investigations are needed to elucidate underlying mechanisms of expediated rod photoreceptor differentiation via 9CRAL.

## ACKNOWLEDGMENTS

We are grateful to Drs. Samuel G. Jacobson and Brian Brooks for skin biopsy samples that were used for generating fibroblasts and subsequently iPSC lines at the Stem Cell Core facility of National Heart, Lung and Blood Institute. We thank Comparative Cytogenetics Core Facility of National Cancer Institute for karyotyping assay. We acknowledge Anupam Mondal, Ben Fadl, Lina Zelinger and Samantha Papal for insightful discussions and constructive comments, and Jacob Nellissery and John Wilson for technical maintenance. This research was supported by Intramural Research Program of the NEI (ZIAEY000450 and ZIAEY000456). Stem cell work in Spain was supported by Spanish Ministry of Science, Innovation and Universities, Instituto de Salud Carlos III (ISCIII)-European Regional Developmental Fund (FEDER) PI16/00409, CP18/00033 and ISCIII-FEDER Platform for Proteomics, Genotyping and Cell Lines, PRB3, PT17/0019/0024. The EM work was funded by FNLCR Contract HHSN261200800001E. The bioinformatic analyses utilized the high-performance computational capabilities of the Biowulf Linux cluster at NIH (http://biowulf.nih.gov)

## Competing Interests

All authors declare no conflict of interest.

## Data availability statement

All raw and processed data are available through Gene Expression Omnibus (www.ncbi.nlm.nih.gov/GEO) with accession GSE129104 and at https://neicommons.nei.nih.gov.

